## Supplementary Figure for "Structure of the catalytically active APOBEC3G bound to a DNA oligonucleotide inhibitor reveals tetrahedral geometry of the transition state"

**Supplementary Data:**

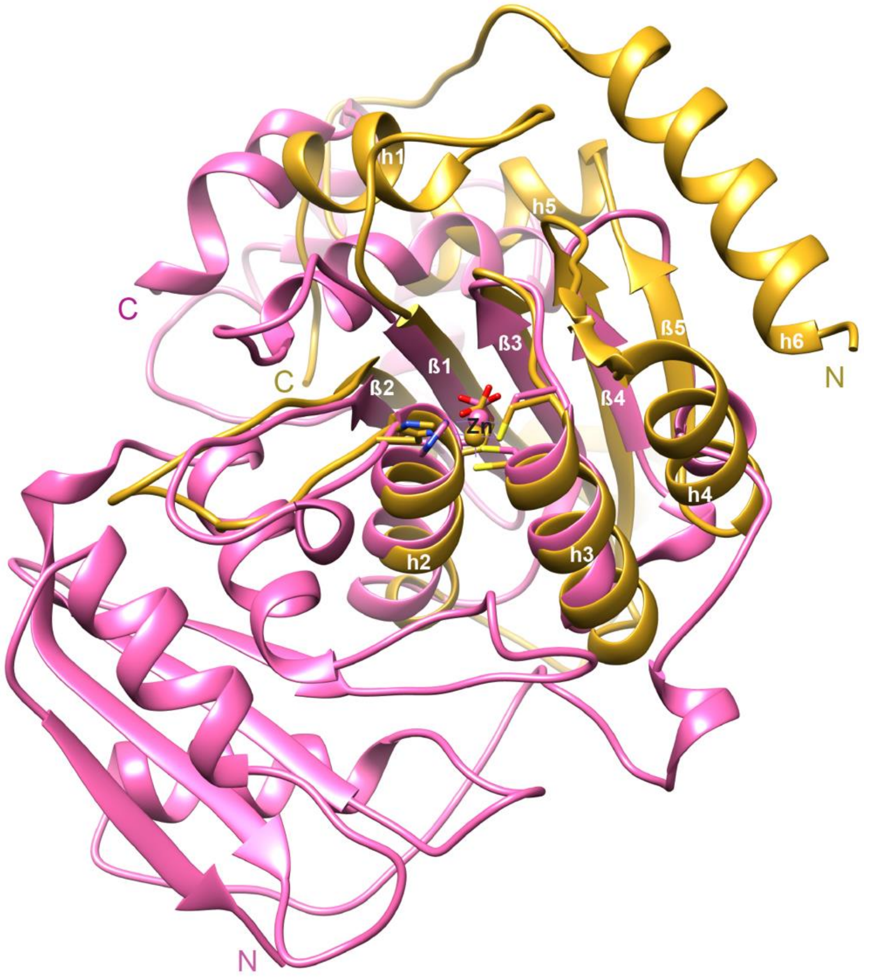


**Supplementary Figure S1a.** Structure alignment of wild-type human A3G-CTD (PDB ID: 4ROV) with *E. coli* CDA (PDB ID: 1CTU) using the program UCSF Chimera. Protein structures are represented as cartoon (A3G-CTD2 is gold and *E. coli* CDA is pink). $Zn^{2+}$ is shown as a sphere (Orange in A3G-CTD2 and pink in *E. coli* CDA). $Zn^{2+}$ coordinating residues (H257, C288, C291 in A3G-CTD and H102, C129, C132 in *E. coli* CDA) and catalytic residue (E259 in A3G-CTD and E104 in *E. coli* CDA) are shown as sticks. N and C (colored as corresponding protein's color) indicate the N- and C-terminal ends of the protein. Only A3G-CTD secondary structures are labeled. $Zn^{2+}$, Helix 2, Helix 3, ß1, ß2, ß3 and ß4 of A3G-CTD are structurally aligned with *E. coli* CDA.

```
A3G-CTD     187 GSHMASLRHSM--------D-PPT-------------------------- 201
E.Coli-CDA    1 -----------MHPRFQTAFAQ--LADNLQSALEPILADKYFPALLTGEQ 37

                                                               ß1
A3G-CTD     202 --------------------FTFNFNNEPWVRGR------------HETY 219
E.Coli-CDA   38 VSSLKSATGLDEDALAFALLP-------------LAAACARTPLSNFNVG 74

                  ß1                    ß2                      h2
A3G-CTD     220 LCYEVERMHNDTWVL-LNQRRGFLCNQAPHKHGF--L---EGRHAELCFL 263
E.Coli-CDA   75 AIARGV---------SG--TWYFGANMEF-----IGATMQQTVHAEQSAI 108

                 h2                  ß3          h3
A3G-CTD     264 DVIP-FWKLDLDQDYR--VTCFTSWSPCFSCAQEMAKFISKNKHVS---L 307
E.Coli-CDA  109 SHAWLSGEK-------ALAAITVNYTPCGHCRQFMNEL---N--S-GLDL 145

                  ß4
A3G-CTD     308 CIFTARIYDDQGRCQEG---------LR--TLAEAGAKISIMTYSEFKHC 346
E.Coli-CDA  146 RIHLP------------GREAHALRDYLPDA------------------- 164

A3G-CTD     347 WDTFVDHQGCPFQPWDGLDEHSQDLSGRLRAILQNQEN------------ 384
E.Coli-CDA  165 --------------------------------------FGPKDLEIKTLL 176

A3G-CTD     384 -------------------------------------------------- 384
E.Coli-CDA  177 MDEQDHGYALTGDALSQAAIAAANRSHMPYSKSPSGVALECKDGRIFSGS 226

A3G-CTD     384 -------------------------------------------------- 384
E.Coli-CDA  227 YAENAAFNPTLPPLQGALILLNLKGYDYPDIQRAVLAEKADAPLIQWDAT 276

A3G-CTD     384 ------------------ 384
E.Coli-CDA  277 SATLKALGCHSIDRVLLA 294
```

**Supplementary Figure S1b.** Structure based sequence alignment of A3G-CTD (PDB ID: 4ROV) with *E. coli* CDA (PDB ID: 1CTU) created by the program UCSF Chimera. Superimposed structures are shown in **Supplemental Fig. S1a**. Structurally aligned sequences (Helix 2, Helix 3, ß1, ß2, ß3 and ß4 of A3G-CTD) are highlighted by yellow. $Zn^{2+}$ coordinating residues (H257, C288, C291 in A3G-CTD and H102, C129, C132 in *E. coli* CDA) and catalytic residue (E259 in A3G-CTD and E104 in *E. coli* CDA) are highlighted by green.

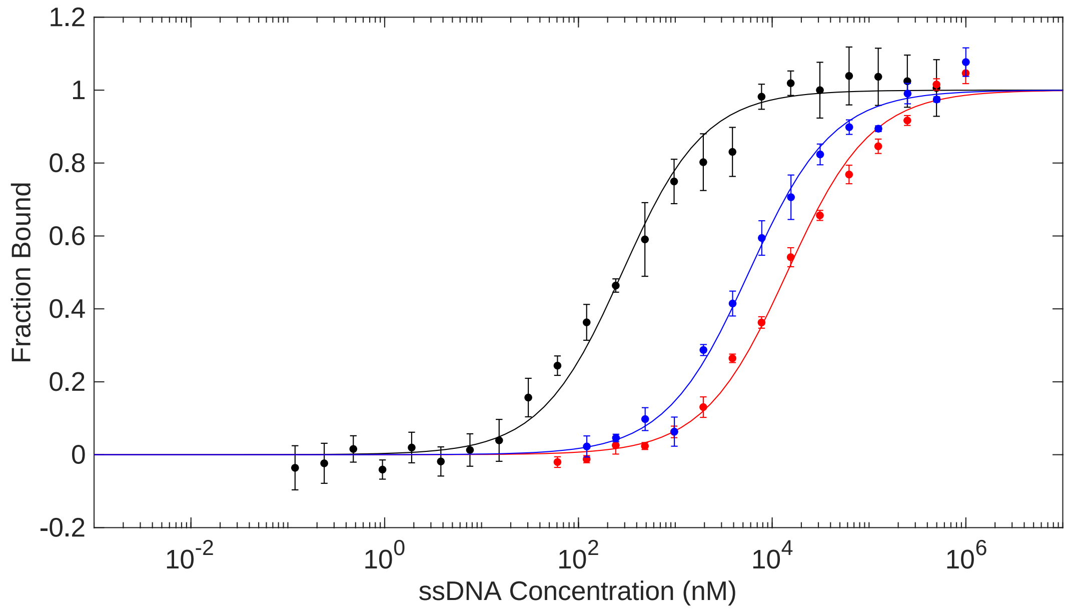


**Supplementary Figure S2a**. Micro Scale Thermophoresis binding data for 5′-AATCCdZAAA binding to active A3G-CTD2 (black), inactive A3G-CTD2* (red), and 5′-AATCCCAAA substrate binding to inactive A3G-CTD2* (blue).

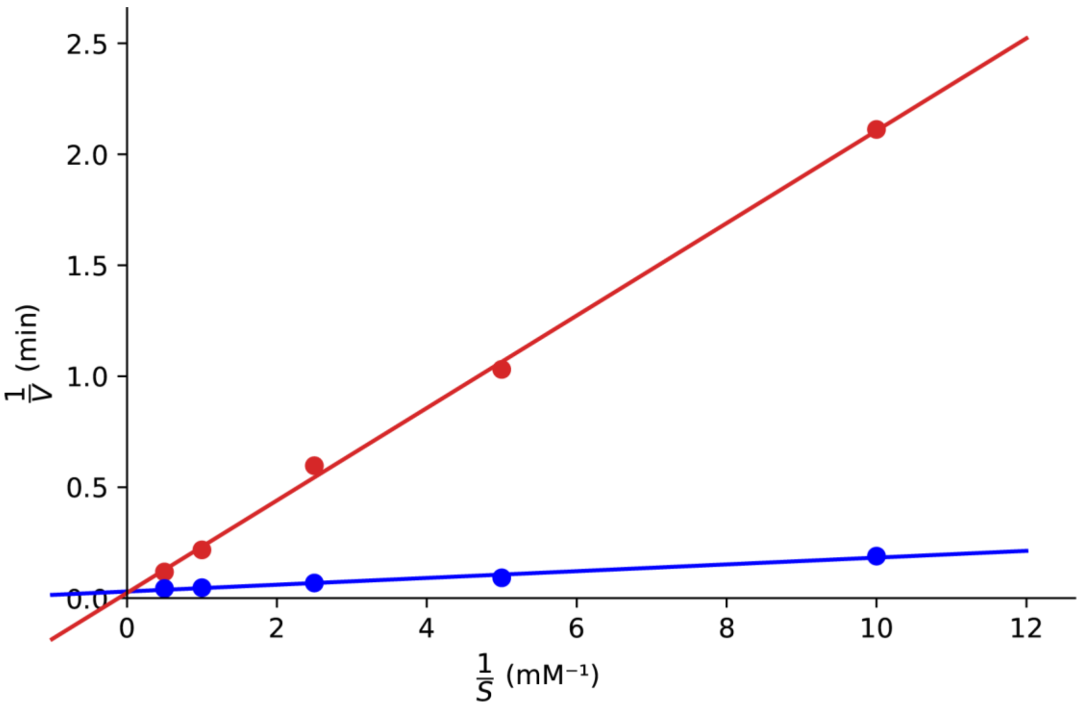


**Supplementary Figure S2b**. Lineweaver Burk plots of 200 nM A3G-CTD2 in the absence (blue) and presence (red) of 50 µM 5′-AATCCdZAAA confirm competitive inhibition of A3G-CTD2 deaminase activity. Initial deamination rates were measured at 100 µM, 200 µM, 400 µM, 1 mM, and 2 mM 5′-AATCCCAAA substrate concentrations.

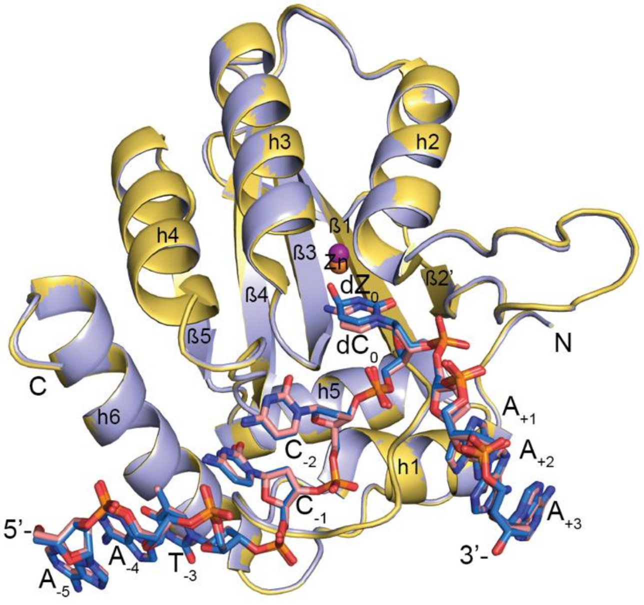


**Supplementary Figure S3.** Superposition of A3G-CTD2:dZ-ssDNA (PDB ID: 7UXD) structure with A3G-CTD2*:ssDNA (PDB ID: 6BUX) structure. Protein structure represented as cartoon (A3G-CTD2 is yellow and A3G-CTD2* is light blue) and ssDNA structure represented as sticks (dZ-ssDNA as blue and dC-ssDNA as pink). $Zn^{2+}$ shown as a sphere (Orange in A3G-CTD2 and purple in A3G-CTD2*). N and C indicate the N- and C-terminal ends of the protein, 5′- and 3′- indicates 5′ and 3′ ends of the ssDNA.

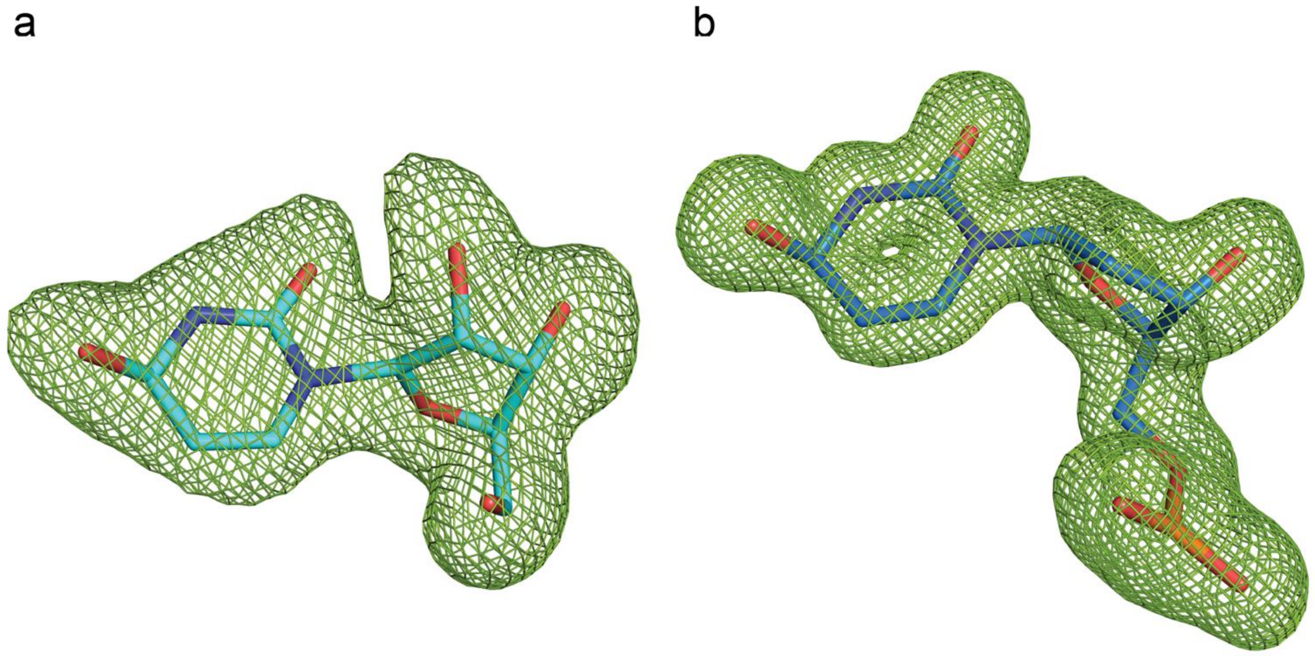


**Supplementary Figure S4.** Omit map for ligands. a) Omit (Fo–Fc) map contoured at 3σ is shown in green around the ribo-zebularine hydrated intermediate (ZEB-OH) (PDB ID: 1CTU)

b) Omit (Fo–Fc) map contoured at 3σ is shown in green around the 2′-deoxy-zebularine hydrated intermediate (dZ-$H_2O$) (PDB ID: 7UXD).

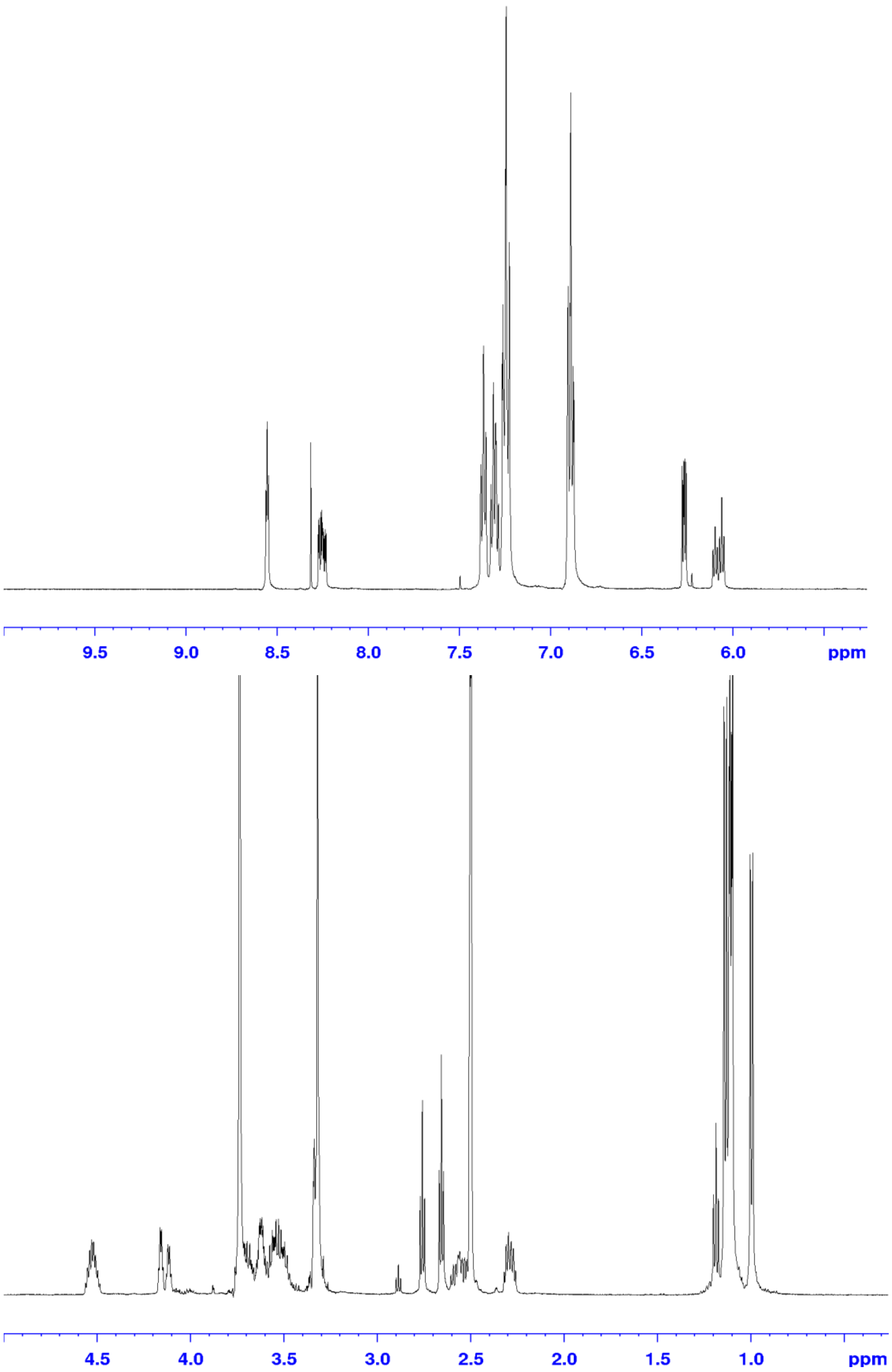


**Supplementary Figure S5.** $^{1}$H NMR spectrum of final dZ phosphoramidite.

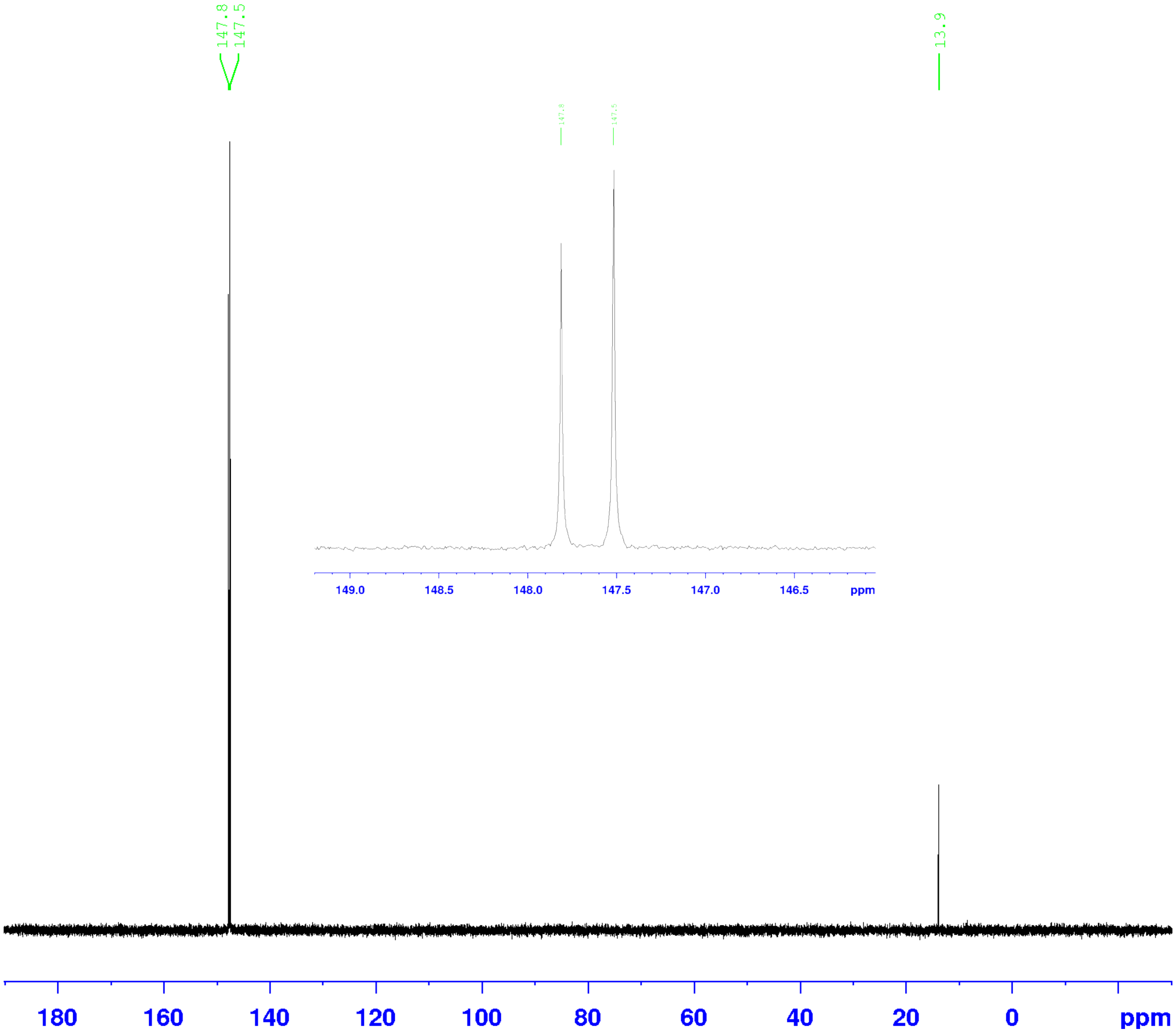


**Supplementary Figure S6.** $^{31}P$ NMR spectrum of final dZ phosphoramidite (inset zoom of 146-149ppm).

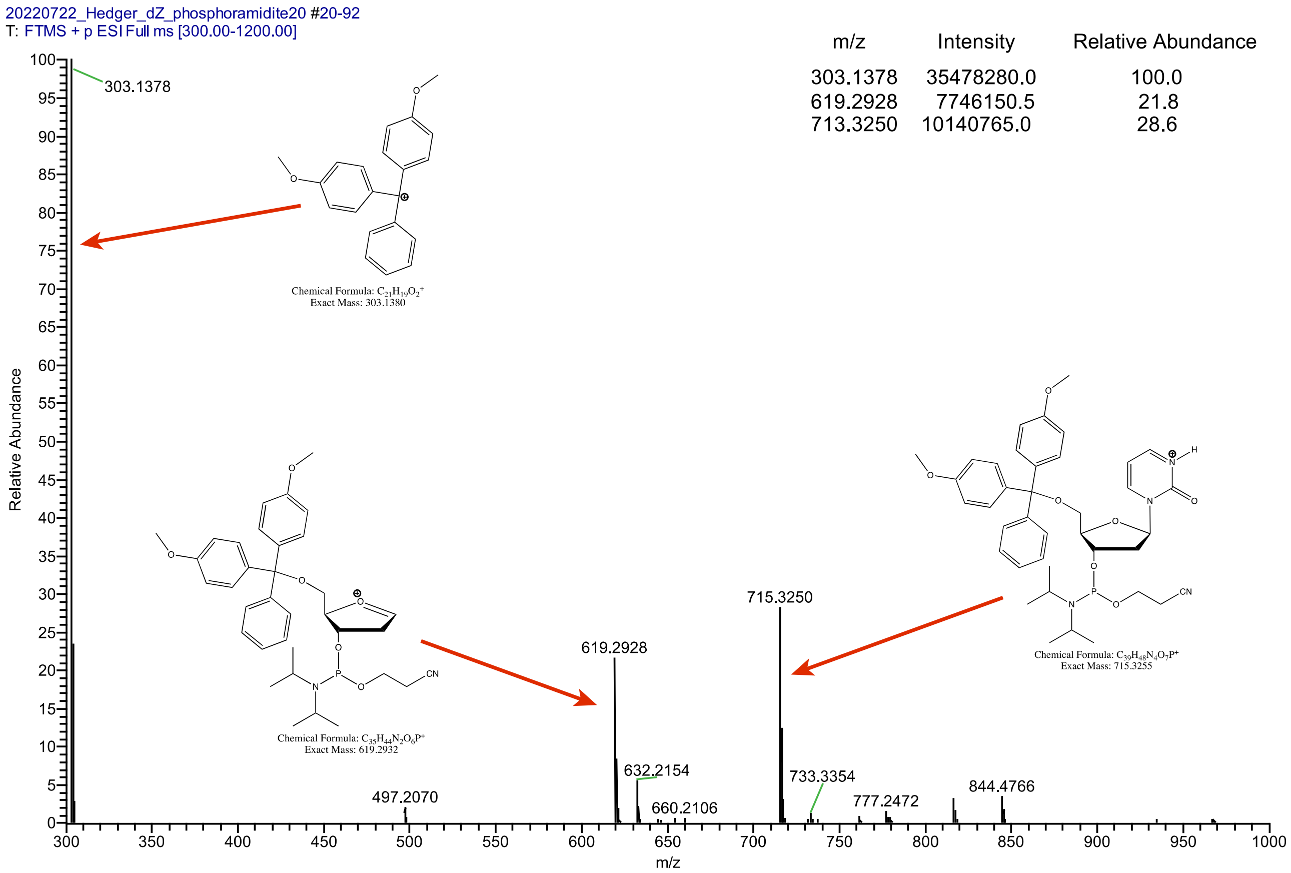


**Supplementary Figure S7.** HRMS ESI Mass spectrum of final dZ phosphoramidite. Spectrum also shows DMT+ cation and oxocarbenium cation fragments from the main compound.

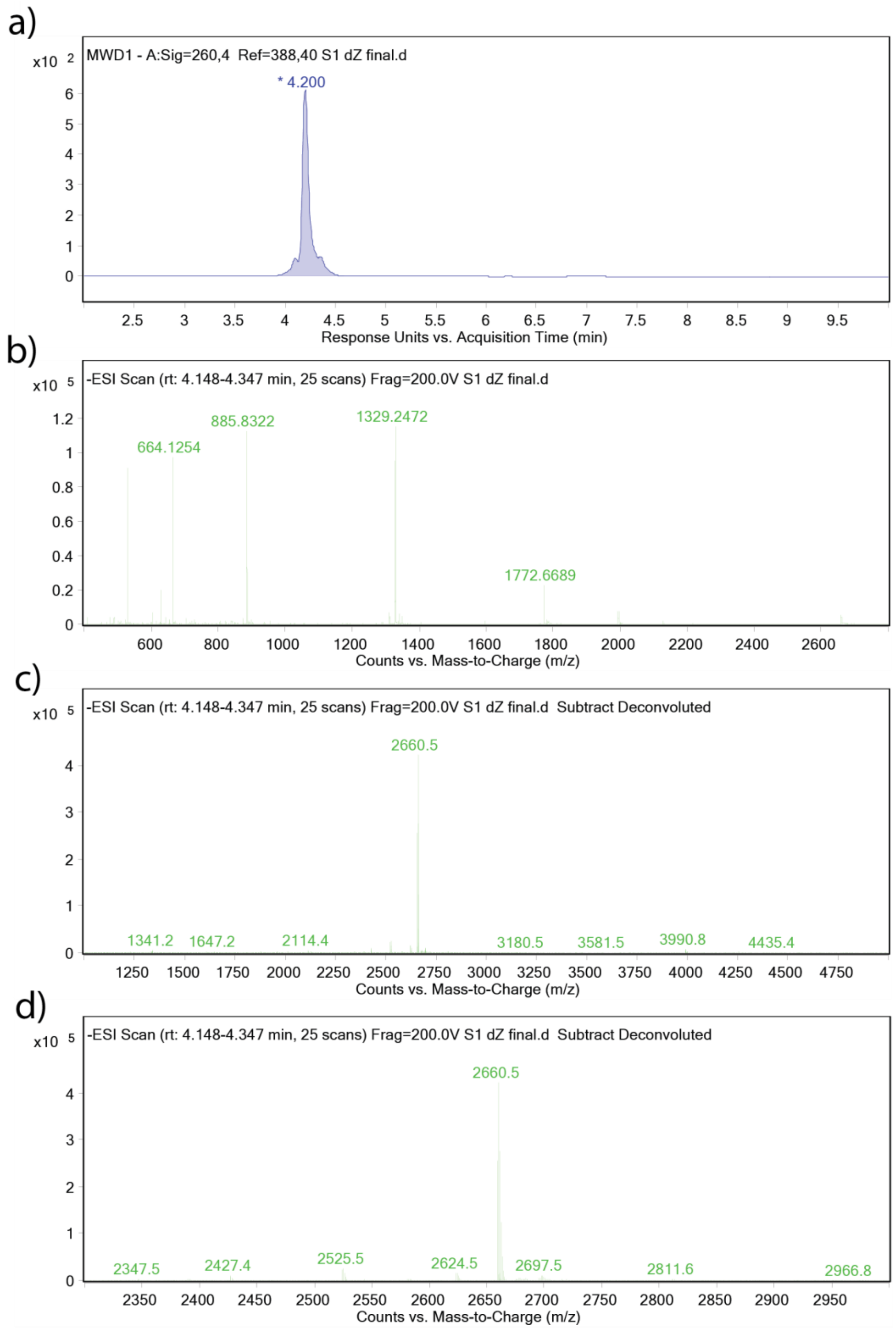


**Supplementary Figure S8.** Mass spectrometry analysis for dZ containing oligonucleotide (5′-AATCCdZAAA). a) LCMS UV trace at 260 nm. b) Mass spectrum showing multiply charged negative species c) Deconvoluted mass spectrum between 1000-5000 m/z showing the main oligonucleotide peak at 2660.5 d) Same as c, but expanded to focus on the region of 2300-3000 m/z.
